# New targets for drug design: Importance of nsp14/nsp10 complex formation for the 3’-5’ exoribonucleolytic activity on SARS-CoV-2

**DOI:** 10.1101/2021.01.07.425745

**Authors:** Margarida Saramago, Cátia Bárria, Vanessa Costa, Caio S. Souza, Sandra C. Viegas, Susana Domingues, Diana Lousa, Cláudio M Soares, Cecília M Arraiano, Rute G. Matos

## Abstract

Severe acute respiratory syndrome coronavirus 2 (SARS-CoV-2) virus has triggered a global pandemic with devastating consequences for health-care and social-economic systems. Thus, the understanding of fundamental aspects of SARS-CoV-2 is of extreme importance.

In this work, we have focused our attention on the viral ribonuclease (RNase) nsp14, since this protein was considered one of the most interferon antagonists from SARS-CoV-2, and affects viral replication. This RNase is a multifunctional protein that harbors two distinct activities, an N-terminal 3’-to-5’ exoribonuclease (ExoN) and a C-terminal N7-methyltransferase (N7-MTase), both with critical roles in coronaviruses life cycle. Namely, SARS-CoV-2 nsp14 ExoN knockout mutants are non-viable, indicating nsp14 as a prominent target for the development of antiviral drugs.

Nsp14 ExoN activity is stimulated through the interaction with the nsp10 protein, which has a pleiotropic function during viral replication. In this study, we have performed the first biochemical characterization of the complex nsp14-nsp10 from SARS-CoV-2. Here we confirm the 3’-5’ exoribonuclease and MTase activities of nsp14 in this new Coronavirus, and the critical role of nsp10 in upregulating the nsp14 ExoN activity *in vitro*. Furthermore, we demonstrate that SARS-CoV-2 nsp14 N7-MTase activity is functionally independent of the ExoN activity. The nsp14 MTase activity also seems to be independent of the presence of nsp10 cofactor, contrarily to nsp14 ExoN.

Until now, there is no available structure for the SARS-CoV-2 nsp14-nsp10 complex. As such, we have modelled the SARS-CoV-2 nsp14-nsp10 complex based on the 3D structure of the complex from SARS-CoV (PDB ID 5C8S). We also have managed to map key nsp10 residues involved in its interaction with nsp14, all of which are also shown to be essential for stimulation of the nsp14 ExoN activity. This reinforces the idea that a stable interaction between nsp10 and nsp14 is strictly required for the nsp14-mediated ExoN activity of SARS-CoV-2, as observed for SARS-CoV.

We have studied the role of conserved DEDD catalytic residues of SARS-CoV-2 nsp14 ExoN. Our results show that motif I of ExoN domain is essential for the nsp14 function contrasting to the functionality of these conserved catalytic residues in SARS-CoV, and in the Middle East respiratory syndrome coronavírus (MERS). The differences here revealed can have important implications regarding the specific pathogenesis of SARS-CoV-2.

The nsp10-nsp14 interface is a recognized attractive target for antivirals against SARS-CoV-2 and other coronaviruses. This work has unravelled a basis for discovering inhibitors targeting the specific amino acids here reported, in order to disrupt the assembly of this complex and interfere with coronaviruses replication.

## Introduction

Over the last years, we have observed the emergence of different coronaviruses (CoVs) that have caused serious human epidemic diseases like the Severe Acute Respiratory Syndrome (SARS) in 2002, the Middle East Respiratory Syndrome (MERS) in 2012 and, currently, the pandemic coronavirus disease-2019 (COVID-19).

SARS-CoV-2 is the causative agent of COVID-19 and belongs, together with SARS-CoV and MERS-CoV, to the genera *Betacoronavirus*. All these three viruses emerged as novel coronaviruses, considered to have initially been transmitted to humans from animals (zoonotic viruses), and all cause respiratory illnesses [1].

CoVs are enveloped, single-stranded positive‐sense RNA viruses from the order *Nidovirales* that have the largest genome among RNA viruses (~30 kb in the case of SARS-CoV-2) [2]. SARS-CoV-2 contains a large replicase gene that occupies two-thirds of the genome encompassing nonstructural proteins (nsps), followed by structural and accessory genes. Among the nsp proteins, there is the nsp14 ribonuclease.

Ribonucleases (RNases) are key factors in the control of all biological processes. These enzymes ensure maturation, degradation and quality control of all types of RNAs in all domains of life [3; 4; 5; 6; 7]. Nsp14 protein has exoribonucleolytic activity conferred by its N-terminal ExoN domain [8]. The ExoN domain resembles the superfamily of DEDDh exonucleases, which also includes the proofreading domains of many DNA polymerases as well as other eukaryotic and prokaryotic exonucleases [9]. These enzymes catalyze the excision of nucleoside monophosphates from nucleic acids in the 3’-to-5’ direction, using a mechanism that depends on two divalent metal ions and a reactive water molecule [10; 11; 12]. This exonucleolytic activity is critical for the proofreading activity during Coronavirus replication, a property missing in other RNA viruses, which enhances its replication fidelity and has played an important role in nidoviral evolution and genome expansion [13].

SARS-CoV-2 and SARS-CoV share 79.5% of genome homology and much of what is presently known about the biology of SARS-CoV-2 was inferred from previous studies on SARS-CoV [14]. However, some striking differences suggest important differences between the two CoVs in terms of infectiousness and the effects they have on human hosts.

Despite the high amino acid sequence identity (95%) between the nsp14 of both viruses, SARS-CoV ExoN knockout mutants are viable, even with an increased mutation frequency, while the equivalent ExoN knockout mutants of SARS-CoV-2 are non-viable [15]. This striking difference suggests an additional and critical ExoN function in SARS-CoV-2 replication. Nsp14 ExoN seems to have a very important role in SARS-CoV-2 RNA synthesis [15], showing up as a prominent target for the development of antiviral drugs.

Whereas basal nsp14 ExoN activity does not require the presence of co-factors, its activity is only fully activated in the presence of the nsp10 protein [16]. The crystal structure of SARS-CoV nsp14 in complex with its nsp10 co-factor has shed light on how the interaction between the proteins occurs [17].

The N7-methylguanosine (m^7^G) cap is a defining structural feature of eukaryotic mRNAs, including those of eukaryotic viruses that replicate in the cytoplasm. SARS-CoV nsp14 was discovered to be a bifunctional protein, since beyond its exoribonucleolytic activity it also displays a guanine-N7-methyltransferase (N7-MTase) activity in its C-terminal domain [18]. This means that the enzyme is capable of methylating cap analogues or GTP substrates, in the presence of S-adenosyl methionine (SAM) as methyl donor [18; 19]. The nsp14 N7-MTtase activity is essential for formation of a functional 5’ RNA cap structure, critical for stability and translation of CoV mRNAs in the host cells. In fact, mRNA cap methylation requires the concerted action of three viral proteins: nsp14, nsp10 and nsp16. Additionally to nsp14, nsp10 is also responsible for stimulating the activity of nsp16 2’-O-MTase, which makes this protein a central player in RNA cap methylation [20]. The obligate sequence of methylation events is initiated by nsp14, which first methylates capped RNA transcripts to generate cap-0 ^7Me^GpppA-RNAs. The latter are then selectively 2’O-methylated by the 2’O-MTase nsp16 in complex with its activator nsp10 to give rise to cap-1 ^7Me^GpppA_2’OMe_-RNAs. While nsp14 recognizes non-methylated RNA cap exclusively, nsp10/nsp16 recognizes N7-methylated RNA cap [20].

Coronaviruses have the inherent capacity to mutate, recombine and infect different hosts. This raises an urgent need for the development of effective antiviral drugs to fight against the present and future pandemic diseases that may arise. Through its dual function, nsp14 protein plays a prominent role in CoV life cycle, and, thus, it is a very attractive target for drug design. Its N7-MTase activity is involved in RNA cap modification to assist in translation and evading host defences, while the 3′-5′ ExoN activity (stimulated by nsp10) has a direct role in CoV RNA synthesis beyond assuring the long-term genome fidelity. In this work we provide a biochemical characterization of SARS-CoV-2 nsp14/nsp10, addressing several aspects of this complex for the first time. By contributing to the deep understanding of nsp14/nsp10 mechanisms of action, this work will also help to clarify its role in the SARS-CoV-2 life cycle. Importantly, it will serve as a basis for the design of effective drugs to treat Covid-19 and other CoVs infections.

## Materials and methods

### Plasmid construction

Full-length nsp10 and nsp14 genes from SARS-CoV2 (Uniprot ID P0DTD1) were optimized for *E. coli* expression and synthesized by GenScript (USA). The synthesized genes were subsequently cloned into the *Nde*I–*Bam*HI sites of commercial pET15b to generate pET15b_Nsp10 and pET15b_Nsp14, which express N-terminal His-tagged versions of nsp10 and nsp14.

The point mutations F19A, G69A, S72A, H80A and Y96A were introduced into pET15b_Nsp10 by overlapping PCR using the primers listed in table S1. The point mutations D90A, E92A, D243A, D273A and D331A were introduced into pET15b_Nsp14 also by overlapping PCR (table S2). All the constructions were verified by sequencing at StabVida (Portugal).

### Protein expression and purification

Plasmids expressing nsp10 WT and point mutations were transformed into Bl21 (DE3) cells, while plasmids expressing nsp14 WT and point mutations were transformed into Rosetta cells for the expression of the recombinant proteins. Cells were grown in LB medium supplemented with 150 μg/ml ampicillin (Nsp10 variants) or TB medium supplemented with 150 μg/ml ampicillin and 50 μg/ml chloramphenicol (nsp14 variants) at 30 °C to an OD600 of 0.5. At this point, protein expression was induced by addition of 0.5 mM IPTG and bacteria were grown for further 16 h. Cells were pelleted by centrifugation and stored at −80°C. To co-purify nsp10 with nsp14, cultures overexpressing each protein separately were pelleted together. The culture pellets were resuspended in 10 ml of Buffer A (40 mM Tris-HCl pH 8, 150 mM NaCl, 10 mM imidazol). Cell suspensions were lysed using the FastPrep-24 (MP Biomedical) at 6.5 m/s for 60 seconds in the presence of 0.5 mM PMSF. The crude extract was treated with Benzonase (Sigma) to degrade the nucleic acids and clarified by a 30 min centrifugation at 10 000 x*g*. Purification was performed in an ÄKTA FPLC™ system (GE Healthcare). The cleared lysate was subjected to a histidine affinity chromatography in a HisTrap HP column (GE Healthcare) equilibrated in Buffer A. Proteins were eluted by a continuous imidazole gradient up to 500 mM in Buffer A. The fractions containing the purified protein were pooled together, and concentrated by centrifugation at 4°C with Amicon Ultra Centrifugal Filter Devices of 10 000 MWCO (Millipore) and buffer exchanged to Buffer B (20 mM Tris-HCl pH 8, 150 mM NaCl). Afterwards, the proteins were subjected to a size exclusion chromatography using a Superdex 200 Increase column (GE Healthcare) with a flow rate of 0.5 ml/min using buffer B. The samples collected were analysed in a 15% SDS-PAGE gel followed by BlueSafe staining (NZYTech, Portugal). Samples with the highest purity were pooled together and concentrated by centrifugation at 4°C with Amicon Ultra Centrifugal Filter Devices of 10 000 MWCO (Millipore). All protein versions were purified at least twice to ensure reproducibility of the results. Proteins were quantified using the Bradford Method and 50% (v/v) glycerol was added to the final fractions prior storage at −20°C.

### RNase Activity Assays

A synthetic 22-mer oligoribonucleotide (H4 5’-UGACGGCCCGGAAAACCGGGCC-3’) (StabVida, Portugal) was used as a substrate in the activity assays. The RNA was labelled at its 5′ end with [^32^P]-γ-ATP and T4 Polynucleotide Kinase (Ambion) in a standard reaction. MicroSpin G-50 columns (GE Healthcare, Cytiva) were used to remove excess [^32^P]-γ-ATP. In order to fold its 3′-end into a duplex structure, the RNA was resuspended in 10 mM of Tris-HCl pH 8.0 and incubated 10 min at 80°C followed by 45 min at 37°C.

The activity assays were performed in a final volume of 12 μl containing the activity buffer - 50 mM HEPES pH 7.4, 1 mM DTT and 5 mM of either MgCl_2_, MnCl_2_, CaCl_2_, NiCl_2_, ZnCl_2_, CoCl_2_ or CuCl_2_ - and the proteins nsp14 and nsp10 (protein concentrations are indicated in the figure legends). The reactions were started by the addition of 50 nM of the RNA substrate, and further incubated at 37°C. Aliquots of 4 μl were withdrawn at the time-points indicated in the respective figures, and the reactions were stopped by the addition of formamide containing dye supplemented with 10 mM EDTA. A control reaction containing only the RNA substrate and the activity buffer (without the enzyme) was incubated in the same conditions during the full time of the assay. Reaction products were resolved in a 20% denaturant polyacrylamide gel (7M urea). Signals were visualized by PhosphorImaging (TLA-5100 Series, Fuji). All the experiments were performed at least in triplicate.

### Surface Plasmon Resonance

Surface plasmon resonance (SPR) was performed by using a CM5 sensor chip (Cytiva) and a Biacore 2000 system (Cytiva). Purified nsp14 was immobilized in flow cell 2 of the CM5 sensor chip by the amine coupling procedure. The surface was activated with a 1:1 mixture of 1-ethyl-3-(3-dimethyllaminopropyl) carbodiimide (EDC) and N-hydroxysuccinimide (NHS), injected during 5 min at a flow rate of 10 μl/min. Then, 20 μg/ml of nsp14 were injected during 10 min at the same flow rate. After the injection of the ligand, ethanolamine was injected over the surface to deactivate it. The immobilization of the protein originated a response of 400 RU. On flow cell 1 (used as a control), BSA protein was immobilized using the same method. Biosensor assays were run at 15 °C in a buffer with 25 mM HEPES-HCl pH 7.4, 0.5 mM DTT and 2,5 mM of MgCl_2_. Serial dilutions of purified Nsp10 WT and mutant proteins were injected over flow cells 2–1 for 3 min at concentrations of 50, 100, 175, 250 and 500 nM using a flow rate of 20 ml/min. The dissociation was allowed to occur during 5 min in the running buffer. All experiments included triple injections of each protein concentration to determine the reproducibility of the signal. Bound proteins were removed after each cycle with a 30 s wash with 2 M NaCl. After each cycle, the signal was stabilized during 1 min before the next protein injection. Data from flow cell 1 was used to correct refractive index changes and nonspecific binding.

### Preparation of the Capped RNA Substrate

A 30-mer RNA substrate was first synthesized using a synthetic DNA template and a promoter oligonucleotide obtained by commercial source (StabVida) for *in vitro* transcription, using the method described by [21]. Briefly, the DNA synthetic template (0.5 μM) and the T7 promoter oligonucleotide (0.6 μM) were annealed in 10 mM of Tris-HCl pH 8.0 by heating for 5 min at 70°C, following by incubation for 30 min, at 37°C. *In vitro* transcription was carried out using ‘NZY T7 High Yield RNA Synthesis kit’ (NZYtech) following manufacturer instructions. To remove the DNA template, 1U of DNase (Invitrogen) was then added to the reaction and incubated 15 min at 37°C. Non-incorporated ribonucleotides were removed with Microspin G-50 Columns (Cytiva).

For the insertion of a ^32^P-labeled cap structure (G*ppp-RNA) in the 5’end of 30 mer RNA substrate we used the vaccinia virus capping enzyme following the manufacturer’s protocol (New England Biolabs), except that the methyl donor SAM was absent and 0.05 units of inorganic pyrophosphatase (New England Biolabs) were added to improve the efficiency of the reaction. A parallel reaction was prepared following the manufacturer’s protocol (New England Biolabs) and using 2 mM SAM to obtain m^7^G*ppp-RNA. Non-incorporated radioisotope was removed using Microspin G-50 Columns (Cytiva), and the labelled substrate was then purified by phenol-chloroform extraction and ethanol precipitation. A small fraction of m^7^G*ppp-RNA and of G*ppp-RNA were digested with 5 μg nuclease P1 (Sigma) in 50 mM NaOAc pH5.2 buffer to originate the m^7^G*ppp and G*pppG markers in the thin-layer chromatography (TLC) run.

### Mtase Activity Assays

To test MTase activity of nsp14 WT or mutant versions, a reaction mix containing the purified recombinant protein (~1 μg), approximately 2 μg of ^32^P-labeled 30-mer G*ppp-RNA substrate, 0.1 mM of SAM, an RNase inhibitor and 2 mM DTT in a total volume of 20 ul, was prepared in the reaction buffer 50 mM HEPES pH 7.4, 1 mM DTT, 5 mM MgCl_2_ and 50 mM KCl, and incubated at 37 °C for 30 min. The same reaction without nsp14 treatment was performed as control. To liberate the cap structures, both RNAs (with and without nsp14 treatment) were digested with 1.25 units of nuclease P1 (Sigma) in 50 mM NaOAc pH5.2 buffer for 30 min at 37°C, followed by the inactivation of the enzyme (75°C for 10 min).

A TLC analysis was performed to separate G*pppG from capped m^7^G*pppG. For this, polyethyleneimine cellulose-F plates (Merck) were previously pre-run with water, air-dried and then spotted with 2-3 ul of the P1 digestion reaction (1 μl spotted at each time, and let dry) and developed in 0.4 M ammonium sulphate. The m^7^G*pppG and G*pppG markers (see above) were run in parallel. The chromatograms were then scanned with a PhosphorImager (TLA-5100 Series, Fuji) to evaluate the extent of ^32^P-labeled capping. All the experiments were performed at least in triplicate.

### Protein modelling

Modeling of the SARS-CoV-2 nsp10/nsp14 complex was done with the program Modeller version 9.23 [22]. The crystal structure of the corresponding complex on SARS-Cov (PDB ID 5C8S) was used as the template for the homology modelling [17]. The catalytic and metal-binding amino acids were kept fixed during the model refinement stages. Twelve independent models were generated and ranked using the normalized DOPE score [23]. Subsequent analyses were carried on the model with the lowest score. The final model was further used as the template for modeling the structures of the mutant proteins. Only the substituted residue and its neighboring residues (within 6 Å) were refined during the modeling process.

## Results

### ExoN activity of nsp14 is stimulated in the presence of nsp10

We have purified both SARS-CoV-2 nsp10 and nsp14 proteins individually through immobilized metal affinity chromatography (IMAC) followed by size-exclusion chromatography based on previous works [16; 24] (Materials and Methods section). As showed in Figure S1, nsp10 and nsp14 migrate at 17 KDa and 62 KDa, respectively, consistent with their expected molecular weights. In the case of nsp10, which is known to have 12 identical subunits assembled into a spherical dodecameric architecture [25], the visible higher bands (highlighted with an asterisk *) correspond to reminiscent oligomers of the protein. This was further confirmed by Western blot using antibodies specific for the His-tag tail of the protein (data not shown).

First, we intended to test the ribonuclease activity of the purified SARS-CoV-2 nsp14. We have used the synthetic RNA substrate H4 previously reported in Bouvet *et al.* [16]. This substrate is a 22-nucleotide (nt) long RNA with a 5’ single-stranded (ss) tail and a 3’-end engaged in a stable duplex structure (Figure S2). The nsp14 ExoN activity was then determined by analyzing the hydrolysis of the H4 RNA 5’-end radiolabeled. As we can see in the left panel of Figure 1A, nsp14 alone exhibits exoribonucleolytic activity; however, in the conditions tested (40 nM of nsp14), the activity was not pronounced. (Figure 1A). Nsp10 was previously reported as a critical co-factor for activation of nsp14 ExoN activity in SARS-CoV [16; 17]. As such, we have analyzed the influence of nsp10 in the activity of SARS-CoV-2 nsp14, using different molar ratios between nsp14 and nsp10 (Figure 1A). SARS-CoV-2 nsp14 ExoN activity was found to be stimulated by nsp10 in a dose-dependent manner. At equimolar ratio (1:1), the ExoN activity is weakly stimulated compared with the ExoN activity exhibited by nsp14 alone. By increasing the concentration of nsp10, the maximal ExoN activity was achieved with a ratio nsp14-nsp10 of 1:4, in agreement with the results reported for SARS-CoV and MERS-CoV [15; 16]. In this condition, we observe a faster RNA degradation and almost complete disappearance of the full-length RNA after 30 minutes of incubation. The degradation products formed range between 17-to 20-nts in length, which corresponds to cleavage from the 3’-end of the RNA duplex region of the 22-nt long H4 RNA. By increasing the concentration of nsp14 (and also of nsp10 proportionally), the hydrolysis of the substrate becomes much more efficient, yielding smaller breakdown products corresponding to degradation of the ss- and double-stranded (ds)-regions (Figure S3). The ladder-like pattern of the degradation products is typical of exoribonucleolytic cleavage of the RNA in the 3’-5’ direction (considering that the RNA is labeled at the 5’ end), further supporting CoVs nsp14 3’-5’ directionality [15; 16; 26]. As a control, we have tested nsp10 alone (at 160 nM, which was the higher concentration used in nsp10:nsp14 ratio) for ribonucleolytic activity under the same conditions used for nsp14. As expected, nsp10 was not able to degrade the RNA substrate (Figure 1A).

**Figure 1.**
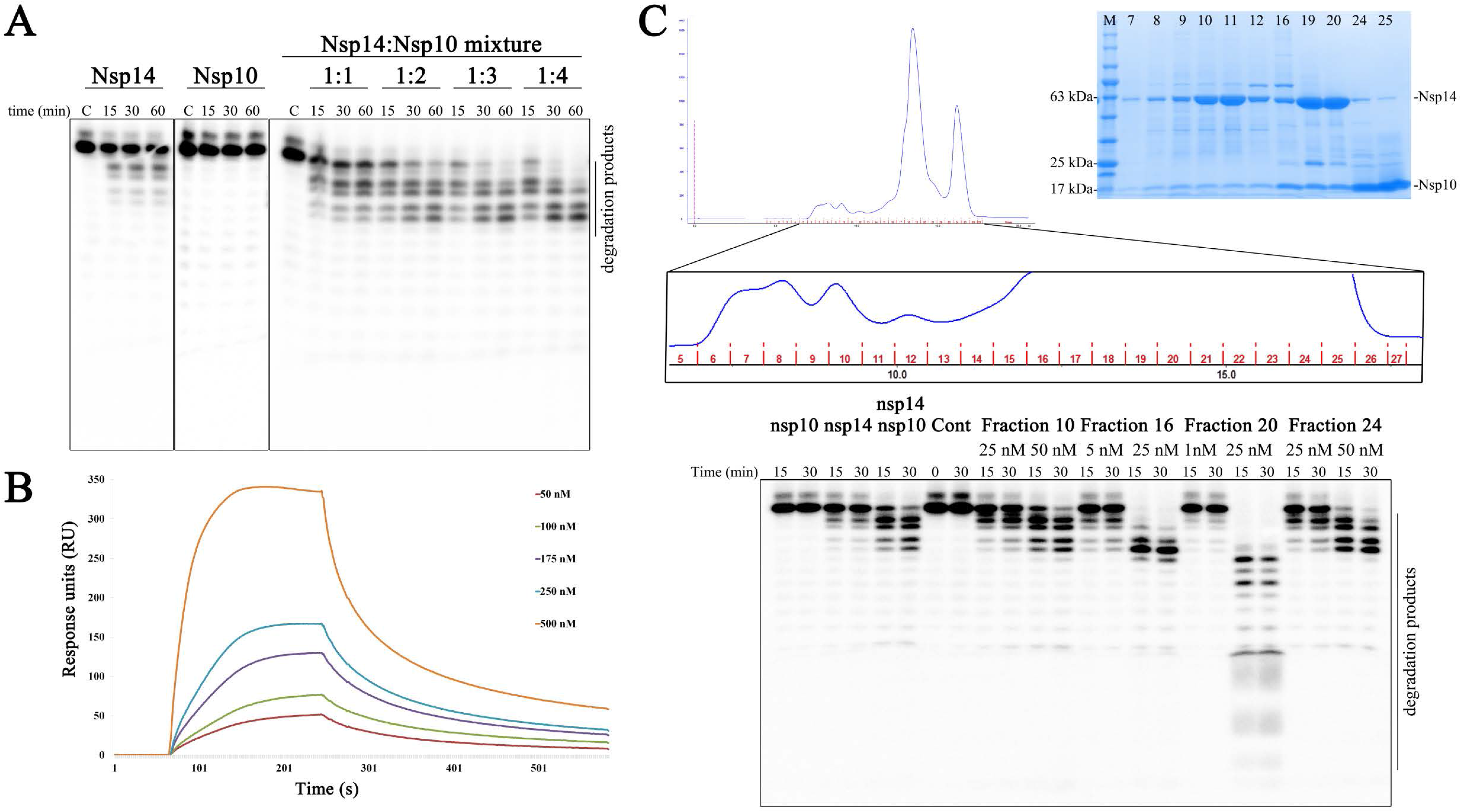
ExoN activity of nsp10:nsp14 complex. **(A)** Activity of nsp14 and nsp10 alone or in combination using 50 nM of H4 RNA substrate. Nsp14 was used at 40 nM in all panels; nsp10 was used at 40 nM in 1:1 ratio, 80 nM in 1:2 ratio, 120 nM in 1:3 ratio and 160 nM in 1:4 ratio and in the second panel from left. **(B)** Surface plasmon resonance analysis. Nsp10 was injected over nsp14 at the concentrations indicated in the figure. The sensorgram represents the average of three independent injections of each concentration. **(C)** Co-purification of nsp14 and nsp10. The chromatogram obtained during the final purification step is presented on the top left. On the top right, a 15% SDS-PAGE gel with the fractions collected during the SEC. In the bottom, the activity of some fractions was analysed using 50 nM of H4 RNA substrate. Protein concentration used is indicated in the figure. Reactions were analyzed on 7 M urea/20% polyacrylamide gels. C, control reactions; time points are indicated in the top of each panel. All the experiments were performed at least in triplicate.

The ability of nsp10 to interact with nsp14 was analyzed through Surface Plasmon Resonance (SPR) (Figure 1B). For this, purified nsp14 was immobilized in a sensor chip and nsp10 was injected at different concentrations as described in materials and methods. The results confirm that both proteins interact with each other as showed by an increased response when nsp10 is injected. This increase is dependent on nsp10 concentrations and it is possible to see the dissociation of nsp10 over time when injection stops. Since nsp14 or nsp10 were shown to be very unstable proteins, it was not possible to obtain a good fit of the data in order to determine the binding kinetics and dissociation constants.

We have also co-purified nsp10 and nsp14 proteins. During the size exclusion chromatography step, several fractions corresponding to different peaks have been collected (Figure 1C). Analysis of those fractions by SDS-PAGE gel shows the presence of both nsp14 and nsp10 proteins in all fractions and in different ratios (Figure 1C), confirming their interaction. We then tested the ExoN activity of some of the eluted fractions (Figure 1C, bottom gel). Fraction 16 that appears to have an equimolar proportion of nsp14-nsp10, exhibited increased activity and degraded the full-length RNA in 15 minutes, with major breakdown products of 18-, 17- and 16-nts. 25 nM of this fraction was sufficient to degrade all the full length substrate, whereas for the mixture of nsp14 and nsp10 separately purified, 40 nM and 160 nM have been used respectively (1:4 ratio). Surprisingly, fraction 20, which appears to have more nsp14 compared to nsp10, was the most active sample. This fraction showed the ability of totally consuming the RNA, starting with the ds-region and then proceeding to the ss-region of the substrate, generating products of smaller size. With 50 nM of this fraction, the degradation proceeds until it reaches a unique and final product of 8-nts (Figure S4). This result is in line with the reported ability of SARS-CoV nsp14 to hydrolyze ssRNA to end products of 8-12 nts [16; 26]. Finally, fractions 10 (excess of nsp14) and 24 (excess of nsp10) present a similar ability to cleave the H4 RNA. The activity of these fractions was comparable to that observed when testing nsp14 and nsp10 mixed together in the 1:4 ratio (Figure 1C). We can conclude that co-purifying nsp14 and nsp10 proteins substantially increases nsp14 ExoN activity, and, in this case, the ratio between them seems not to be determinant to achieve high levels of ExoN activity. However, taking into account that our intention was the study of several mutant versions of these proteins from SARS-CoV-2, and considering the results obtained, the remaining experiments were performed with nsp14 and nsp10 purified separately. The fact that both nsp14 and nsp10 need a ratio in which nsp10 is in excess (Figure 1A), may be an indication of the instability of the complex, otherwise it might have maximum activity with a 1:1 ratio. Indeed, during the experimental part of this work, we noticed that the ExoN activity decreased with time and the activity is completely lost after two weeks. The same behavior was previously observed [27]. As such, all the experiments were performed with proteins freshly purified.

### SARS-CoV-2 nsp14 ExoN activity is metal-dependent

The activity of the nsp14 ExoN domain is also known to be dependent on divalent cations [16; 17; 26; 28]. In order to determine which divalent metals supports maximal activity of the SARS-CoV-2 nsp14-nsp10 complex, we tested ExoN activity in the presence of different divalent ions: Mg^2+^, Mn^2+^, Ca^2+^, Ni^2+^, Cu^2+^, Co^2+^ or Zn^2+^ (Figure 2). As already reported for several other RNases, binding of these different metal ions might affect the activity of SARS-CoV-2 ExoN by inducing structural changes in the active site [28; 29; 30]. The results show that SARS-CoV-2 ExoN nsp14 is active in the presence of both Mg^2+^ and Mn^2+^, with a more pronounced activity in the presence of Mg^2+^. When Mn^2+^ is added to the reaction, the activity is more distributive and smaller breakdown products with ladder-like pattern are visible (Figure 2). Ca^2+^ did not support the catalysis, in agreement with data obtained for SARS-CoV nsp14 and other proteins from the DEDD family [11; 31]. The same was observed for Ni^2+^ and Cu^2+^ ions (Figure 2). Interestingly, in the presence of Co^2+^ and Zn^2+^ we observed a residual, but not relevant ExoN activity (Figure 2). Chen *et al*. [28] also reported nsp14 residual activity in the presence of Zn^2+^. Finally, the addition of the chelating agent EDTA to the reaction completely blocks nsp14 ExoN activity (Figure 2), confirming the importance of divalent ions, namely Mg^2+^ and Mn^2+^, for this activity, similar to that described for the SARS-CoV counterpart [16; 27; 28].

**Figure 2.**
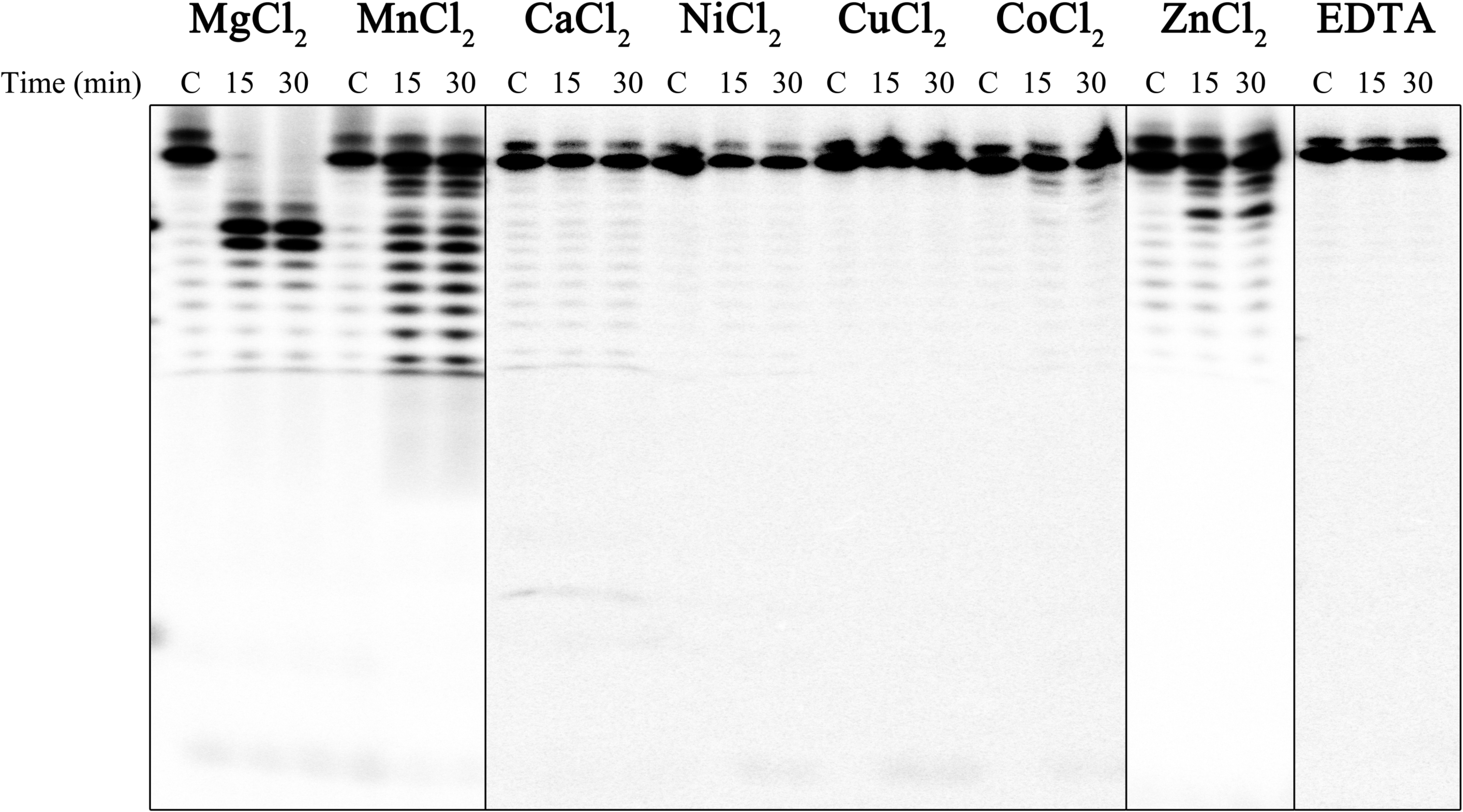
Metal cofactor-dependent activity of nsp14. nsp14:nsp10 complex (40 nM:160 nM) was incubated with 5 mM of different divalent ions. Reactions were analyzed on 7 M urea/20% polyacrylamide gels. C, control reactions; time points are indicated in the top of each panel. All the experiments were performed at least in triplicate.

### SARS-CoV-2 nsp10 residues involved in nsp14-10 complex formation showed that the stability of this complex is determinant for ExoN activity

Until now, there is no available structure for the SARS-CoV-2 nsp14-nsp10 complex. As such, we have modelled the SARS-CoV-2 nsp14-nsp10 complex based on the 3D structure of the complex from SARS-CoV (PDB ID 5C8S) (Figure 3) [17]. This was possible due to the sequence and structural similarities between nsp10 and nsp14 proteins between the two viruses (Figure 4A and 5A). The crystal structure of the isolated nsp10 from SARS-CoV-2 was recently solved, and all-atoms RMSD between this structure and the one from the SARS-CoV nsp10-nsp14 complex is 1.36 Å [32]. However, the isolated nsp10 structure from SARS-CoV-2 was not used in the modelling, since we wanted to keep the nsp14-nsp10 native contacts as much as possible. The all-atoms RMSD between the generated model (Figure 3) and the crystal structure of SARS-CoV nsp10-nsp14 complex was 0.76 Å. Comparing with the SARS-CoV-2 nsp10 structure, our model showed an all-atoms RMSD of 1.33 Å, which is in agreement with the deviations found between the crystal structures. The model is represented in Figure 3, and as we can see the interaction between nsp14 and nsp10 is figuratively similar to a “hand (nsp14) over fist (nsp10)”.

**Figure 3.**
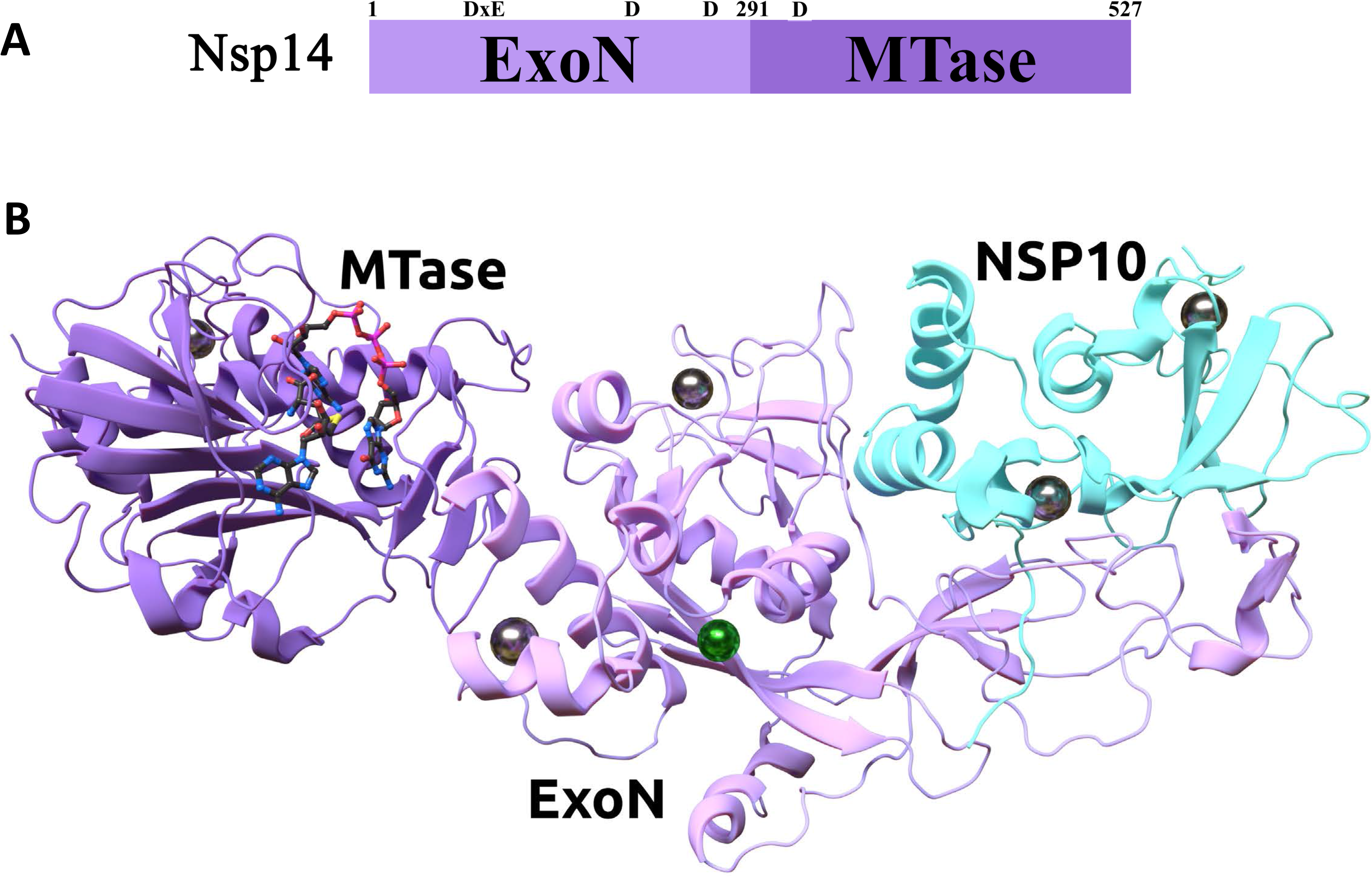
**(A)** Scheme of nsp14 domain organization. ExoN domain, aa 1-291; MTase domain, aa 292-527. **(B)** Homology model of the nsp10-nsp14 complex of SARS-CoV-2. The nsp10 is marked in cyan. The MTase and ExoN domains of nsp14 are marked in dark and light purple, respectively. The complex was refined in the presence of Mg^2+^ (green sphere) and Zn^2+^ (gray spheres) ions, and the substrates SAH and GpppA (sticks).

**Figure 4.**
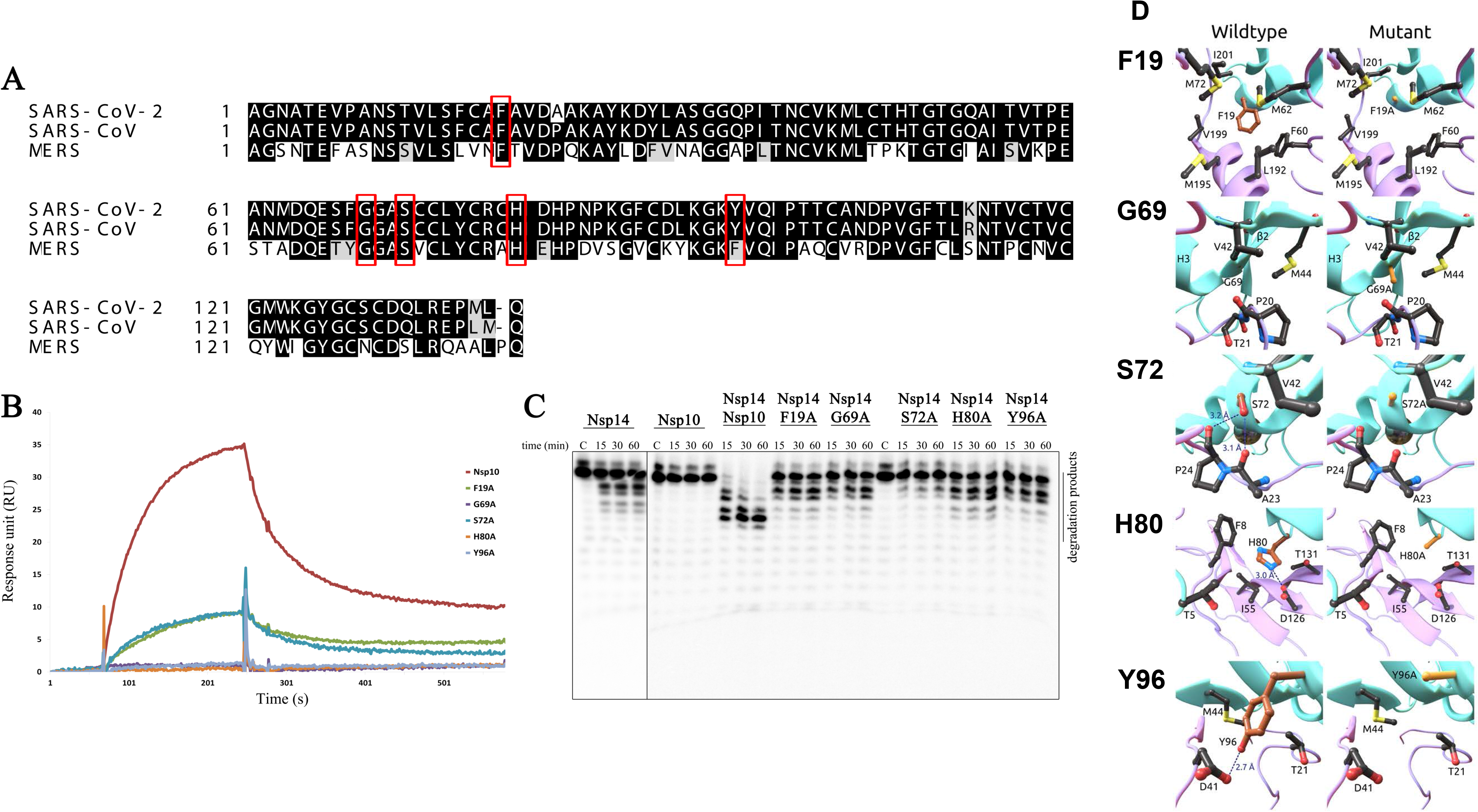
Influence of nsp10 mutations on nsp14-nsp10 complex formation and ExoN activity. **(A)** Sequence alignment of nsp10 from SARS-CoV-2 (Uniprot ID: P0DTD1), SARS-Cov (Uniprot ID: P0C6×7) and MERS (YP_009047225). Residues mutated in this work are highlighted with a red box **(B)** Surface plasmon resonance analysis. Nsp10 WT and mutant versions were injected over nsp14 at a concentration of 75 nM. The sensorgram represents the average of three independent injections of each protein. **(C)** Activity of nsp14 (40 nM) in the presence of nsp10 WT and mutant versions (160 nM). Reactions were analyzed on 7 M urea/20% polyacrylamide gels. C, control reactions; time points are indicated in the top of each panel. All the experiments were performed at least in triplicate. **(D)** Position of the nsp10 mutations and their neighboring residues. The WT and mutated structures of F19A, G69A, S72A, H80A and Y96A mutations are depicted on the left and right columns, respectively. Carbons of the WT and mutated residues are colored in brown and orange, respectively. Hydrogen bond and salt-bridges are marked in dashed lines with the respective distances. The nsp10 and nsp14 secondary structures are marked in cyan and purple, respectively.

**Figure 5.**
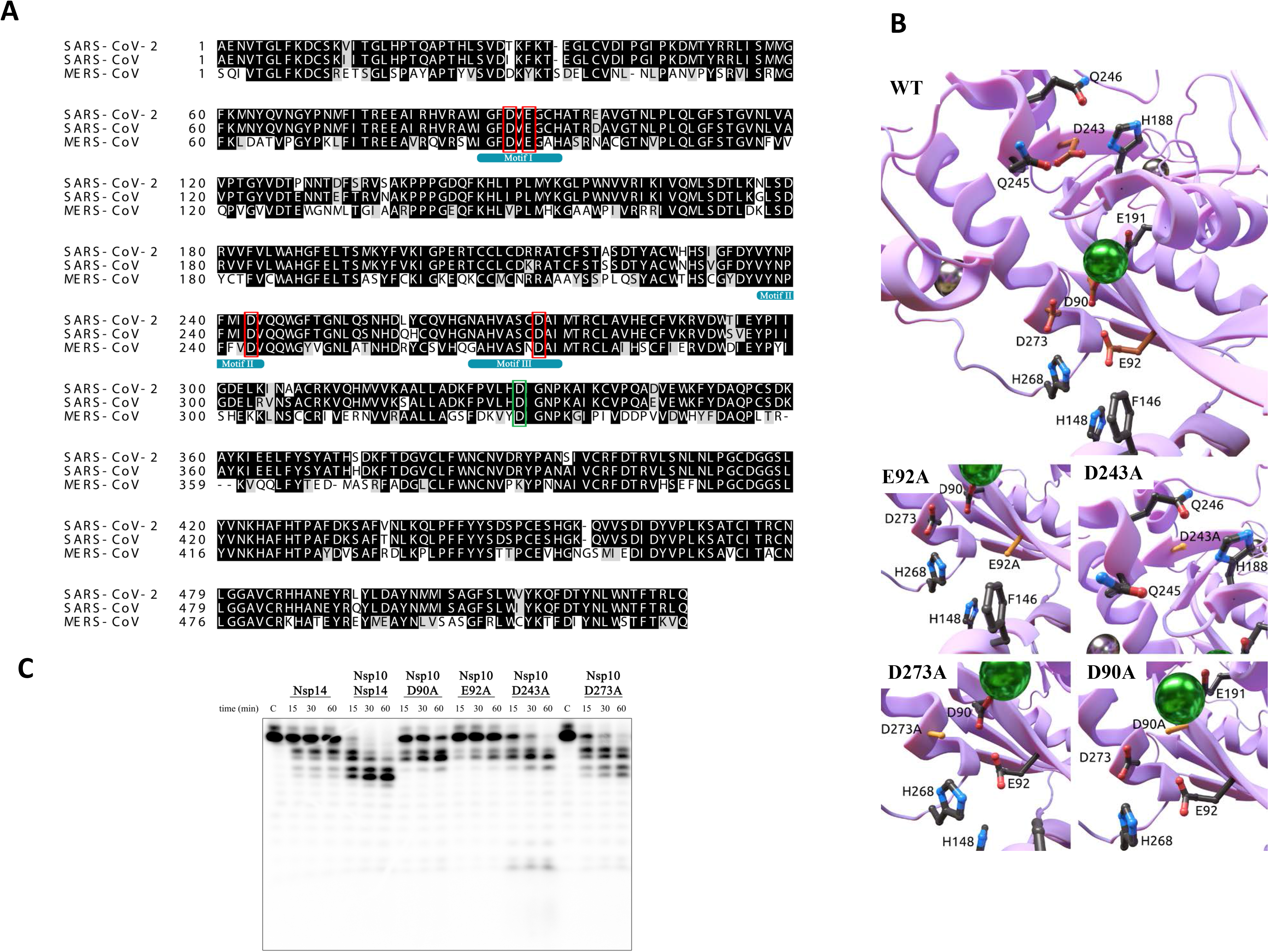
Analysis of DEDD motif mutations for ExoN activity. **(A)** Sequence alignment of nsp14 from SARS-CoV-2 (Uniprot ID: P0DTD1), SARS-CoV (Uniprot ID: P0C6×7) and MERS (YP_009047225). Residues from the DEDD motif are highlighted with a red box and D331 in a green box. **(B)** Structure of the modeled nsp14 ExoN domain and respective mutants. Overall structure of the WT ExoN domain with the side chain of residues subjected to alanine substitutions (carbon atoms marked in brown) and other residues that may participate in the exoribonuclease reaction. Magnesium and Zinc ions are depicted in green and gray spheres respectively. The structure of the mutations E92A, D243A, D273A and D90A are shown with carbons colored in orange. The side chains of non-mutated residues were kept fixed during the structure refinement. **(C)** Activity of nsp14 WT and mutant versions (40 nM) in the presence of nsp10 WT (160 nM). Reactions were analyzed on 7 M urea/20% polyacrylamide gels. C, control reactions; time points are indicated in the top of each panel. All the experiments were performed at least in triplicate.

Conserved amino acids located on the surface of nsp10 were reported to be involved in nsp14-nsp10 interaction in other CoVs [16; 24; 33; 34]. As such, and in order to study their role in ExoN activity, we mutated some of these amino acids (highlighted in red boxes in Figure 4A) into alanines creating the point mutants F19A, G69A, S72A, H80A and Y96A. As shown in the sequence alignment of the SARS-CoV-2 nsp10 protein with the nsp10 from the two other highly pathogenic and deadly human coronaviruses SARS-CoV and MERS (Figure 4A), these amino acids are conserved among the three CoVs with the exception of Y96. This residue is unique to SARS-CoV and SARS-CoV-2, whereas a phenylalanine (F96) is found in most other CoVs, including MERS that belongs to a different lineage of the *betacoronavirus* genus [34]. Indeed, the sequence alignment (Figure 4A) reveals that the three proteins present a high degree of sequence conservation, with SARS-CoV-2 nsp10 being more closely related to SARS-CoV (97.1% of sequence identity and 99.3% of sequence similarity).

Substitution of these residues by an alanine is not expected to drastically alter the structures of these nsp10 protein variants comparing to that of the nsp10 wild-type (WT), as confirmed by NMR for SARS-CoV nsp10 mutants [33; 34; 35]. His-tagged nsp10 mutant derivatives were expressed in *E. coli* and purified (Figure S1). The functional consequences of these nsp10 mutations were evaluated in terms of their interaction with nsp14 through SPR. As expected the instability of these proteins, only enabled us to access the interaction of 75 nM of each version with immobilized nsp14, without determining the kinetics of the reaction. As we can see in Figure 4B, F19A and S72A have their ability to interact with nsp14 reduced, whereas G69A, H80A and Y96A have apparently completely lost their ability to interact with nsp14 WT in these conditions.

According to our homology model represented in Figure 4D, the alanine substitutions affect the binding between nsp10 and nsp14 in different ways. The F19A mutation seems to weaken the van der Waals interactions of nsp10 with the helix H4 of the nsp14 (Figure 4D). This can represent the loss of an important hydrophobic effect at the interface of the two proteins. On one end, this helix is part of the framework that keeps key catalytic residues and an Mg^2+^ ion in place in the ExoN active site, whereas the other end is extended by a loop that forms a zinc finger that is crucial for the structural stability of the nsp14 from SARS-CoV [17]. The binding of nsp10 to this region avoids the exposure of hydrophobic amino acids to the solvent and helps the stabilization of the protein structure. The residue G69 is located in a known structural motif that links the nsp10 sheet β2 to the helix H3 (Figure 4D). This motif is part of an extensive region of intermolecular interactions between nsp10 and nsp14 that also helps on the stabilization of the ExoN domain [16]. The presence of a glycine amino acid in this position also allows for the accommodation of the carbonyl group of nsp14 P20. In this scenario, the mutation G69A is likely to cause local changes to the nsp10 secondary structure and to create unfavorable interactions with P20 and, thus, may interfere on the fitting between nsp10 and nsp14. This same region contains the residues S72, H80 and Y96, which participate in the network of polar interactions that connects nsp10 to nsp14. Specifically, residue H80 forms a salt-bridge with nsp14 D126 (Figure 4D) while residue Y96 keeps a hydrogen bond with the carboxyl group of nsp14 D141 (Figure 4D). Residue S72 is a special case as it makes hydrogen bonds with the carbonyl of nsp14 residues A23 and P24 (Figure 4D). Thus, the substitution of these residues by alanine breaks intra and intermolecular interactions that may be important not only for the binding between nsp10 and nsp14 but also to maintain their secondary structure.

Taken together, these mutations on nsp10 are prone to interfere on the interaction of this protein with nsp14. Evidence points to a stabilization role of nsp10 on the nsp14 structure and, thus, mutations that alter the binding between these proteins may disrupt the structure of the nsp14 ExoN domain and decrease its activity [16; 17].

Since any alteration on the binding ability of nsp10 to nsp14 might modify the capacity of nsp14 ExoN to efficiently cleave RNA, we have also analyzed the effect of these nsp10 mutations on the activity of SARS-CoV-2 nsp14 ExoN. Time-course *in vitro* assays with 5’-end labelled H4 RNA were performed in the presence of nsp10 WT or mutants (Figure 4C), using the 1:4 nsp14-nsp10 ratio as described above. All the nsp10 substitutions tested severely affected the nsp14 ExoN activity. Full disappearance of the RNA substrate was not detected even 60 minutes after incubation with any of the nsp10 variants, contrarily to that verified with the nsp10 WT, which allowed total degradation of the full-length RNA in 15 min. The major degradation products obtained from these reactions range between 19 and 20 nts, which correspond to cleavage in the ds-region of H4 RNA, observed for nsp14 alone, and no further progression is observed (Figure 4C). The most striking result was obtained with the S72A derivative that not only completely lost the ability to stimulate the exoribonucleolytic activity of nsp14, but partially compromise the basal ExoN activity of nsp14.

Overall, our results show that all nsp10 mutants have reduced nsp14 affinity when compared to the WT protein, with consequences for the stimulation effect on nsp14 ExoN activity (Figures 4B and C). In agreement with this, the nsp10 F19A, G69A, S72A, H80A and Y96A mutants were also shown to affect nsp14-nsp10 interaction in SARS-CoV, and resulted in a low nsp14 ExoN activity, suggesting an important role for these residues in nsp14 recognition and interaction [24]. These results reinforce the idea that a stable interaction between nsp10 and nsp14 is strictly required for the nsp14-mediated ExoN activity of SARS-CoV-2.

### ExoN motif I has a prominent role on the RNase activity of SARS-CoV-2 nsp14

SARS-CoV-2 nsp14 also has a high similarity with nsp14 from SARS-CoV (95% of sequence identity and 99.1% of sequence similarity), and less conservation with the amino acid sequence of MERS (Figure 5A). As a bimodular protein, the amino acids 1– 290 from SARS-CoV-2 nsp14 fold into the ExoN domain, and the amino acids 291–527 form the N7-MTase domain (Figures 3). In nsp14 the differences are spread punctually over the protein structure. However, none of these positions are inside the MTase or the ExoN active sites. The ExoN activity depends on the conserved DEEDh motif, which is part of the active site (Figure 5A) [13; 16; 26]. These conserved active site residues were found to be distributed over three canonical motifs (I, II, and III). The SARS-CoV-2 residues D90/E92 (motif I), D243 (motif II) and D273 (motif III) are fully conserved in the three CoVs represented. Amino acid residues E92, D243 and D273 form an electrophilic environment that is important for substrate binding and catalysis on nsp14 of SARS-CoV-2 (Figure 5B). Additionally, E92 is probably responsible for the coordination of the second Mg^2+^ ion necessary for the reaction. It may also form a hydrogen bond with H268, which is another key residue that anchors the substrates. We have constructed and purified nsp14 mutants with single substitutions of these conserved catalytic residues by the neutrally charged alanine (D90A, E92A, D243A and D273A) (Figure S1). We conducted activity assays using these variants in order to assess the role of these amino acids in the ExoN activity of SARS-CoV-2 nsp14. The SARS-CoV and MERS nsp14 studies available so far reported that substitution of ExoN catalytic residues by alanine have a large impact in their exoribonucleolytic activity [15; 16; 17; 26]. As shown in Figure 5C, mutations on these residues influenced the ExoN activity of SARS-CoV-2 nsp14. E92A substitution had the most striking effect, presenting only residual exoribonuclease activity. Although to a less extent, the D90A mutant was also severely deficient in their ability to degrade RNA and rendered a major product of 19 nts. These results demonstrate the importance of residues from motif I (D90 and E92) for the activity of nsp14. In contrast, mutations in aspartates from motifs II and III (D243A and D273A, respectively) led to a modest reduction in the ExoN activity, both being able to fully degrade the full-length RNA after 60 min of incubation (Figure 5C). However, the cleavage patterns generated with both mutants were different. The nsp14 D273A version yielded products ranging between 17- and 20-nts (more similar to that of WT), whereas the D243A mutant generated major products between 18- and 20-nts. However, this mutant version was able to go further in the degradation since smaller minor products could be detected (Figure 5C).

Due to the extra role of the motif I E92 residue, it is likely that the substitution E92A causes a major impact on the catalysis compared to the mutations D243A and D273A. The residue D90 also coordinates an Mg^2+^ ion (Figure 5A), but surprisingly the impact of the mutation D90A was not as severe as the observed on E92A. Probably the presence of the residue E191 may be sufficient to keep the Mg^2+^ ion in the active site, attenuating the effects of the D90A substitution. This behavior is consistent with the data from SARS-CoV that shows that the E191 mutation is more likely to affect the ExoN activity than the mutation D90A [17].

In general, our observations contrast with the results obtained for SARS-CoV, and recently for MERS [15; 16; 17; 26]. In the closely related SARS-CoV, the frequently used motif I-double substitution D90A/E92A resulted in a major reduction of ExoN activity, whereas the motif II D243A and motif III D273A mutations completely abrogated the ExoN activity [16; 17; 26]. Also in MERS, both D90A and E92A substitutions appeared to be slightly less detrimental than D273A, even though all of them resulted in almost complete loss of ExoN activity [15]. The results here reported revealed some differences on the functionality of the conserved catalytic residues D90, E92E, D243 and D273 from nsp14 ExoN, which can be related with the pathogenesis of SARS-CoV-2.

### Nsp14 N7-MTase Activity is functionally independent of the ExoN activity

As already mentioned, nsp14 is a bifunctional enzyme with both ExoN and N7-MTase activities, connected by a hinge region that modulates the flexibility of the protein (Figure 3). The C-terminal MTase domain of nsp14 was found to be able to methylate cap analogues or GTP substrates, in the presence of S-adenosylmethionine (SAM) as methyl donor [18; 36]. In this work, we have established an *in vitro* assay with purified SARS-CoV-2 nsp14 to test its N7-MTase activity, using capped 30-mer G*ppp-RNA as substrate (Figure 6). An nsp14 D331A mutant was constructed and used as a negative control due to its reported involvement in the binding of the methyl donor S-adenosyl-methionine (SAM) in SARS-CoV [16; 18; 36]. The level of cap methyltransferase activity of nsp14 was evaluated by the conversion of G*pppRNA to m^7^G*pppRNA during the cap methyltransferase reaction in TLC analysis. Subsequent P1 digestion of the RNA substrates to individual nucleotides gives the ratio of m^7^G*pppG spot compared to G*pppG on the TLC plate phosphor image. SARS-CoV-2 nsp14 was able to methylate the G*pppG cap of RNAs since we can observe, in the corresponding lane, a main spot that co-migrates with m^7^G*pppG marker (Figure 6A). In contrast, treatment of G*pppRNA with nsp14 D331A mutant did not generate a similar product on the TLC plate (Figure 6A). Instead, it gives a nucleotide product that co-migrates with G*pppG marker resultant from the digestion of G*pppRNA with P1 nuclease.

**Figure 6.**
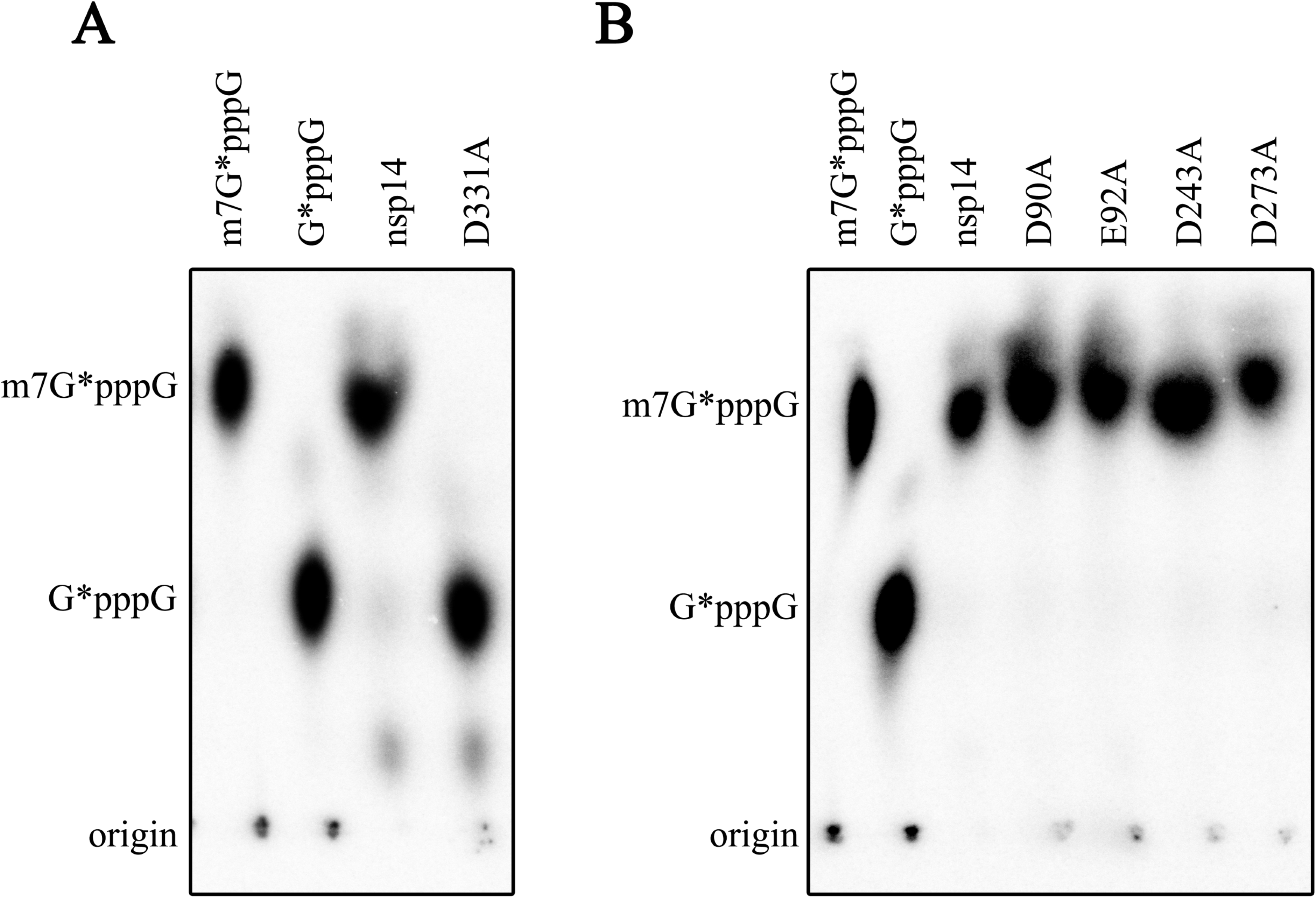
Nsp14 MTase activity. **(A)** TLC analysis of nuclease P1-resistant cap structures released from the G*ppp-RNA methylated by nsp14 WT and D331A mutant. **(B)** TLC analysis of nuclease P1-resistant cap structures released from the G*ppp-RNA methylated by nsp14 WT and nsp14 ExoN catalytic mutants D90A, E92A, D243A and D273A. On the left of both images, we have the P1 digestions of the m^7^G*pppRNA and G*pppRNA produced by the commercially available vaccinia capping enzyme, that were used as markers for m^7^G* and G*. The positions of origin and migration of m^7^G*pppG/G*pppG (lanes 1–2) are indicated on the right.

To evaluate if the nsp14 ExoN activity has any influence on the SARS-CoV-2 MTase activity, the point mutations located in the conserved motifs of the ExoN domain (D90A, E92A, D243A and D273A) were also tested for their capacity to methylate the G*pppRNA in the presence of SAM. As shown in Figure 6B, none of these substitutions influence nsp14 MTase activity, in line with [36].

It has been reported that nsp10 exclusively interacts with the ExoN domain of nsp14 without perturbing the N7-MTase activity [16; 37]. In fact, our results demonstrate that nsp14 presents a robust MTase activity in the absence of nsp10, similar to the one observed with the nsp14-nsp10 complex (data not shown).

From this part of the work we can conclude that the C-terminal region of SARS-CoV-2 nsp14 functions as an MTase, and that this domain is functionally independent of the D90A, E92A, D243A and D273A ExoN catalytic residues. The nsp14 MTase activity also seems to be independent on the presence of nsp10 cofactor, contrarily to nsp14 ExoN.

## Discussion

The 16 non-structural proteins (nsp1-16) encoded by the coronavirus genome (ORF1a/1b) are involved in viral replication and represent potential targets for antiviral drug discovery. Among these, nsp14 is a bifunctional enzyme that harbors both N-terminal ExoN and C-terminal N7-MTase activities [17; 18; 26]. The combination of both activities is unique for coronaviruses. Its ExoN activity is stimulated through the interaction with other non-structural protein, nsp10, which constitutes a critical regulator of viral RNA synthesis and degradation. Due to the central role of the nsp14-nsp10 complex, we have performed a detailed characterization of these proteins from SARS-CoV-2, the coronavirus responsible for the current COVID-19 pandemic.

Nsp10 residues involved in the interaction with nsp14 were found to be essential for SARS-CoV and MHV replication [24; 33]. This reflects the importance of the nsp10-nsp14 interaction surface in coronaviruses. Our results indicate that the same is valid for SARS-CoV-2, since nsp14 only shows a weak ExoN activity in the absence of nsp10, which is strongly enhanced upon interaction with nsp10 (also observed in [27]). We have further established that the concentration ratio for the nsp10/nsp14 complex of SARS-CoV-2 is 4:1, as this yielded maximal ExoN activity. The same ratio was observed for SARS-CoV [16].

Although the 3D structure of the SARS-CoV-2 nsp14-nsp10 complex is not yet solved, our work gave the first insights into important SARS-CoV-2 nsp10 residues directly involved in the interaction with nsp14. This is here demonstrated by the SPR data, as inferred by the homology model done on the basis of the SARS-CoV experimentally determined structure, and the high sequence identity between the proteins from the two viruses. We demonstrate that mutations in the nsp10 F19, G69, S72, H80 and Y96 amino acids residues have deleterious consequences for the nsp14 ExoN activity *in vitro*, which could be rationalized by the structural effects that these mutations have in the formation of the nsp10-nsp14 heterodimer. Therefore, we infer that a stable interaction between these two proteins is strictly required for the correct functioning of nsp14 ExoN activity of SARS-CoV-2. Nsp10 mutations that disrupted the nsp10-nsp14 interaction were lethal for MHV and SARS-CoV, but the inactivation of the ExoN domain was not. However, this domain was found to be essential for the viability of SARS-CoV-2 (and also MERS) [15; 24; 33; 38; 39]. Thus, the nsp14 ExoN activity might have a predominant role on SARS-CoV-2 life-cycle.The role of nsp10 is reinforced by its pleiotropic function during viral replication. Additionally to nsp14, nsp10 is also responsible for stimulating the nsp16 2’-O-MTase activity [20]. Nsp14 and nsp16 share a common interaction surface with nsp10. Some residues here identified as crucial for SARS-CoV-2 ExoN activation, have already been recognised as essential for nsp16 2′-O-MTase activity [16; 24; 40]. Y96 targets both the nsp10-nsp14 and nsp10-nsp16 interactions in SARS-CoV [24]. The same seems to be valid for SARS-CoV-2 based in our results *in vitro* and on the crystal structure of the nsp10-nsp16 complex recently solved [41]. An alanine substitution in the residue S72 of nsp10, which has also been reported to be involved in SARS-CoV nsp10-nsp16 and nsp10-nsp14 interactions [24], resulted in a total loss of SARS-CoV-2 nsp10 ability to stimulate the exoribonucleolytic activity of nsp14. This mutant even seems to compromise the nsp14 basal activity, usually observed in the absence of nsp10. Therefore, inhibition of SARS-CoV-2 nsp10 could have an impact on several steps of viral RNA synthesis.

Our results highlighted nsp10 as a potential target for antiviral drug development. We propose some nsp10 residues that can be targeted to disrupt both the nsp14-nsp10 and the nsp16-nsp10 complexes, leading simultaneously to inhibition of ExoN and 2′-O-MTase activities. Because this protein is highly conserved among CoVs, molecules developed to inhibit SARS-CoV-2 nsp10 interaction surface might be extended to other coronaviruses. To support this, Ogando and colleages [15] reported the interchangeability of nsp10 between SARS-CoV and MERS. Indeed, the nsp10-derived peptide TP29, which was developed to target MHV nsp16 2’-O-MTase activity, successfully suppressed SARS-CoV replication in cell culture [42]. Importantly, nsp10 may be targeted by inhibitors with minimum cross-reactivity with human proteins, since no structures with a fold similar to nsp10 were found in prokaryotes or eukaryotes [32].

We have also looked into the role of conserved DEDD catalytic residues for the ExoN activity of SARS-CoV-2 nsp14. Our results revealed that both motif I D90 and E92 amino acids are strictly required for the RNase activity of this protein, whereas mutations on motif II D243 and motif III D273 do not play such an important role. Thus, we would expect that D90 and E92 residues may have a large impact on SARS-CoV-2 replication. All the nsp14 D90A/E92A, D243A and 273A substitutions that impaired SARS-CoV ExoN activity *in vitro*, also displayed drastic effects *in vivo*. Less accumulation of viral RNA, defects in the synthesis of subgenomic RNAs, and a failure to recover infectious viral progeny was observed in HCoV 229E [26]. The same mutations abrogated detectable RNA synthesis and gave rise to nonviable MERS-CoV [15].

The most explored role of nsp14 ExoN so far is the repairing of mismatches that may be introduced during CoVs RNA synthesis. This capability has also been proposed to be responsible for the excision of nucleoside analogs that are incorporated into RNA, and lead to premature termination of viral RNA replication and survival [38; 39; 43; 44]. Thus, the inhibition of nsp14 ExoN proofreading may potentiate the effect of nucleoside-based inhibitors, such as Remdesivir and Favipiravir [45; 46]. It was previously seen that SARS-CoV and MHV mutants lacking nsp14 ExoN activity exhibited increased susceptibility to nucleoside inhibitors [24; 38; 39; 43; 44]. In particular, an MHV deficient in ExoN proofreading was significantly more sensitive to Remdesivir [45]. For these reasons, the combination of nucleoside analogs with nsp14 inhibitors may be more effective. As such, our biochemical results may be very important for the design of new molecules to inhibit the ExoN activity from SARS-CoV-2, which should be targeting the two important residues for catalysis, D90 and E92. These indications, together with the lack of sequence homology with the human proteome [47], makes nsp14 an excellent druggable protein.

Pathogenic viruses that replicate in the cytoplasm have evolved mechanisms to facilitate infection of mammalian cells. These include the generation of cap structures on their RNA through methyltransferases (MTases) [48]. In this study, we have demonstrated for the first time the nsp14 N7-MTase activity of SARS-CoV-2. This activity is a key factor for equipping viral mRNAs with a functional 5’-terminal cap structure in order to be recognized by the cellular translation machinery. Unmethylated capped RNAs can induce antiviral innate immune responses [49; 50]. Here, we have reported that the nsp14 D331A substitution affects nsp14 MTase activity, which lost the capacity to methylate a capped RNA transcript to generate cap-0 ^7Me^GpppA-RNAs. This makes this residue also of special interest as a target of putative drugs.

Besides viral life cycle, nsp14 can also influence the immune response of the host cells. During the replication process of CoVs, the generated dsRNA intermediates are known to activate the type I interferon (IFN-I) response [51]. Nsp14 was indeed identified as one of the most potent interferon antagonists from SARS-CoV-2 [52]. In the present work, our results revealed the ability of nsp14 ExoN to cleave dsRNA. Thus, this protein may be involved in the degradation of viral dsRNA replication intermediates, hindering the activation of the host innate immune response. This hypothesis was already pointed for SARS-CoV and MERS-CoV [15; 16].

As reported for other CoVs, our results also confirm that the 3’-5’ exoribonucleolytic activity of nsp14 is dependent on metal ions, preferentially Mg^2+^ over Mn^2+^, Co^2+^ and Zn^2+^ promote residual activity, whereas Ca^2+^, Ni^2+^ and Cu^2+^ did not support catalysis. It is tempting to speculate on divalent metals availability as major environmental determinants of the RNase activity of nsp14 *in vivo*. In this regard, alteration of ion homeostasis in favor of infection has been already demonstrated in several viral systems, including in SARS-CoV and MERS [53]. Even trace metals such as zinc and copper are known to influence the course and the outcome of a variety of viral infections [54]. More experimentation and structure information is nonetheless required to fully understand the precise functional and structural role of individual residues in SARS-CoV-2 nsp10 and nsp14 proteins. The determination of the 3D structure of the nsp10/nsp14 complex will be also helpful to increase our knowledge about the synergies between these proteins and to validate our findings. Also, the effect of the nsp14 and nsp10 mutations here described should be employed in a reverse genetic approach in SARS-CoV-2 to study viral replication and transcription.

It was already demonstrated that CoVs ExoN activity of nsp14, which is stimulated by nsp10, constitutes a critical regulator of viral RNA synthesis and degradation. Our results pinpoint the residues that are crucial for the complex formation and ExoN activity, which may be very important to discover new inhibitors that may be used to treat COVID-19 and other diseases caused by other CoVs.

## Acknowledgements

We are grateful to the distinguished and experienced virologists Miguel Fevereiro and Margarida Henriques (INIAV, Lisbon, Portugal) for the valuable discussions regarding physiology of SARS-CoV-2. We also thank Teresa Batista da Silva for technical support.

This work was funded by national funds through FCT - Fundação para a Ciência e a Tecnologia, I. P., Project MOSTMICRO-ITQB with refs UIDB/04612/2020 and UIDP/04612/2020. Project PTDC/BIA-BQM/28479/2017 to R.G.M, and project PTDC/CCI-BIO/28200/2017 to D.L. R.G.M and was also financed by an FCT contract (ref. CEECIND/02065/2017). SCV was financed by FCT program IF (ref. IF/00217/2015). M.S., S.D. and D.L. were financed by an FCT contract according to DL57/2016, [SFRH/BPD/109464/2015], [SFRH/BPD/84080/2012] and [SFRH/BPD/92537/2013], respectively. V.C. and C.B. have a fellowship and a contract, respectively, under the project PTDC/BIA-BQM/28479/2017.C.S.S. was financed by a fellowship under the project ShikiFactory100, grant agreement number 814408 from the European Union’s Horizon 2020 research and innovation programme.

## Supplementary Material

### 1.1. Supplementary Figures

**Figure S1.**
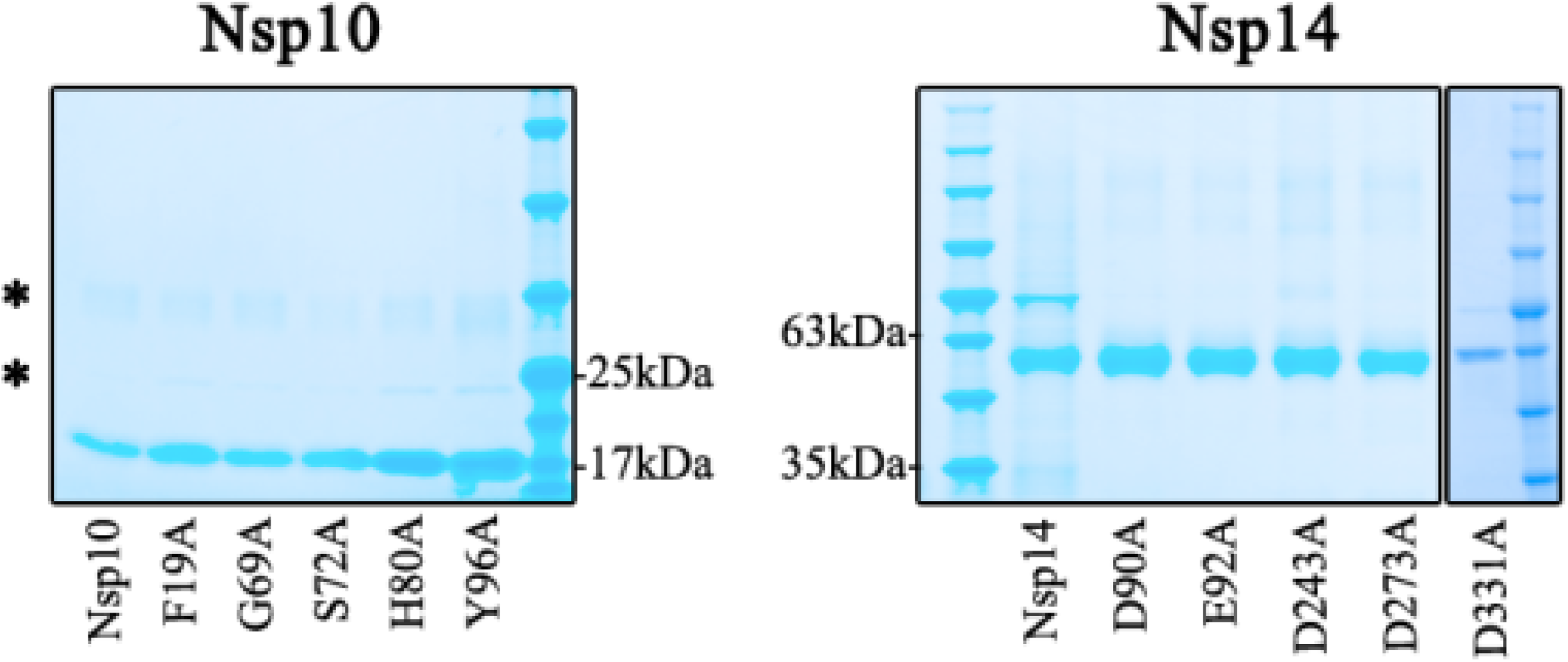
Purification of SARS-CoV-2 nsp10 and nsp14 WT and mutant proteins. SDS-PAGE analysis of the purified nsp10 WT and mutant versions F19A, G69A, S72A, H80A, Y96A (gel in the left), and nsp14 WT and respective mutant versions D90A, E92A, D243A, D273A, D331A (gel in the right). Samples were denatured and separated in a Novex™ 8-16% Tris-Glycine Gel (Invitrogen™). Gel was stained with BlueSafe (NZYTech, Portugal) to visualize protein bands. NZYColour Protein Marker II (NZYTech, Portugal) was used as a molecular weight marker and its respective band sizes are represented in both gels.

**Figure S2.**
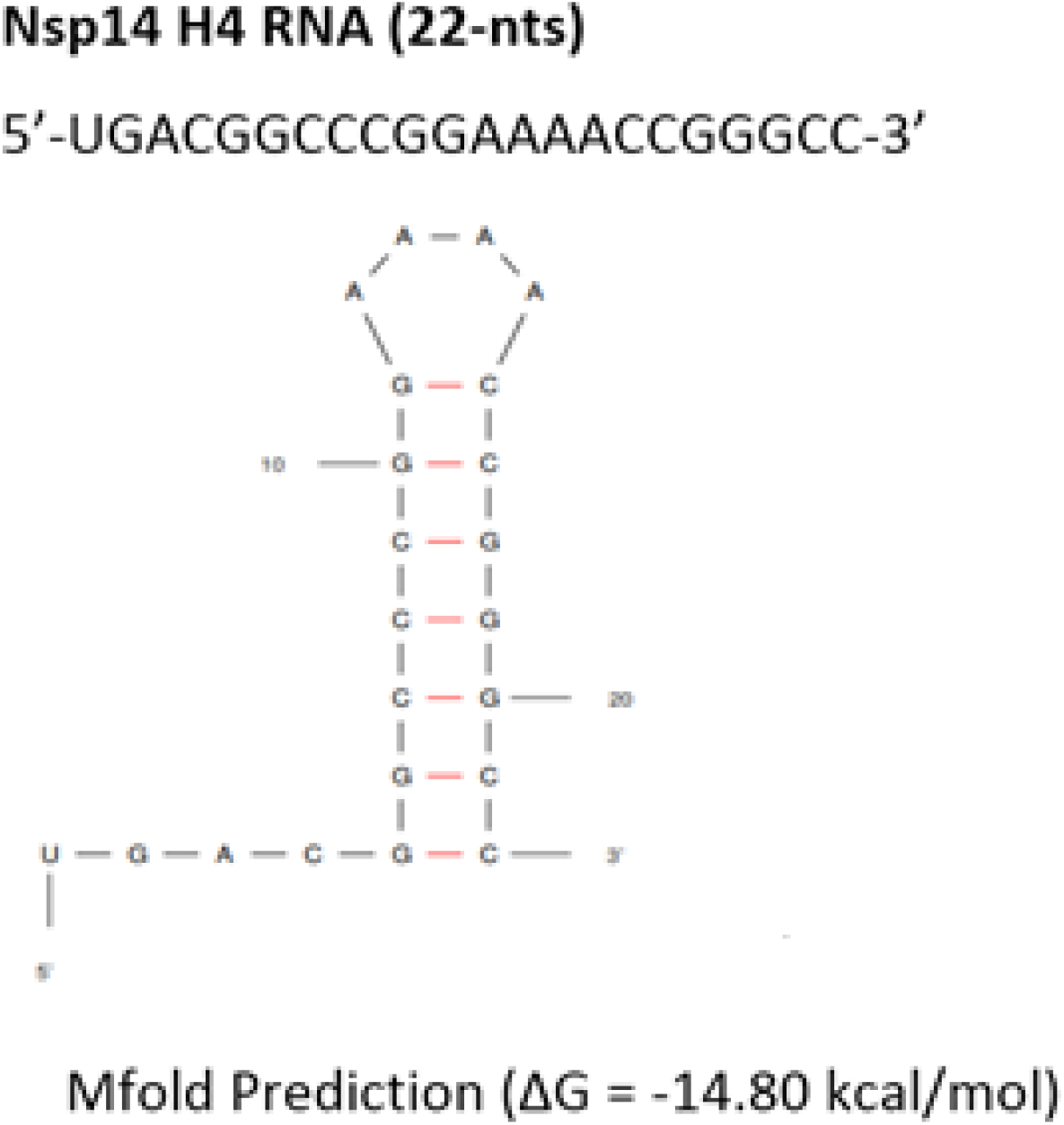
Predicted structure adopted by H4 RNA substrate used in the RNase assays. The H4 RNA sequence, its most stable predicted secondary structure obtained using Mfold RNA modelling server (http://www.unafold.org/RNA_form.php) and the respective minimum free energy (ΔG) are represented [1].

**Figure S3.**
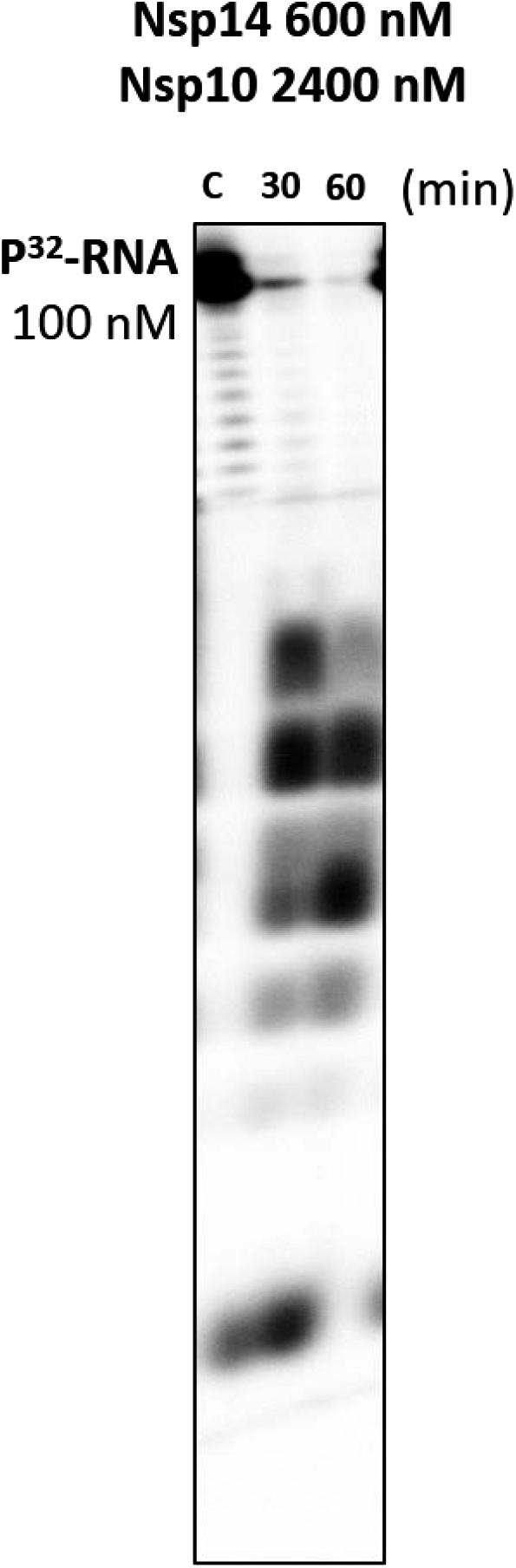
ExoN activity of nsp14 in the presence of nsp10. The activity of nsp14 WT (600 nM) in the presence of nsp10 WT (2.400 nM) was analyzed using 100 nM of H4 RNA substrate. Reactions were run on 7 M urea/20% polyacrylamide gel. C, control reactions; time points are indicated in the top of each panel.

**Figure S4.**
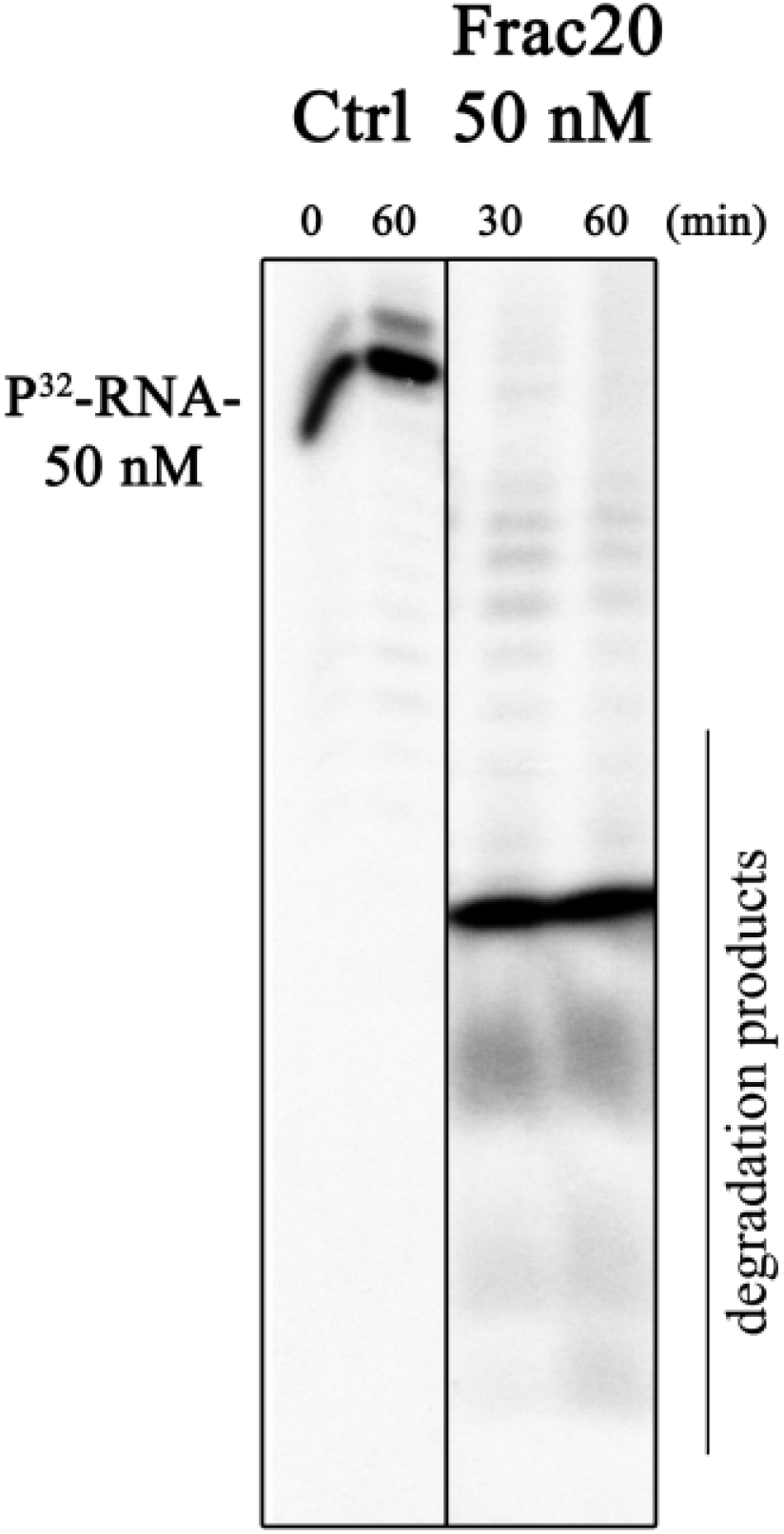
ExoN activity of nsp10:nsp14 complex. The activity of fraction 20 (50 nM) was analyzed using 50 nM of H4 RNA substrate. Reactions were run on 7 M urea/20% polyacrylamide gel. C, control reactions; time points are indicated in the top of each panel.

### 1.2. Supplementary Tables

**Table S1.**
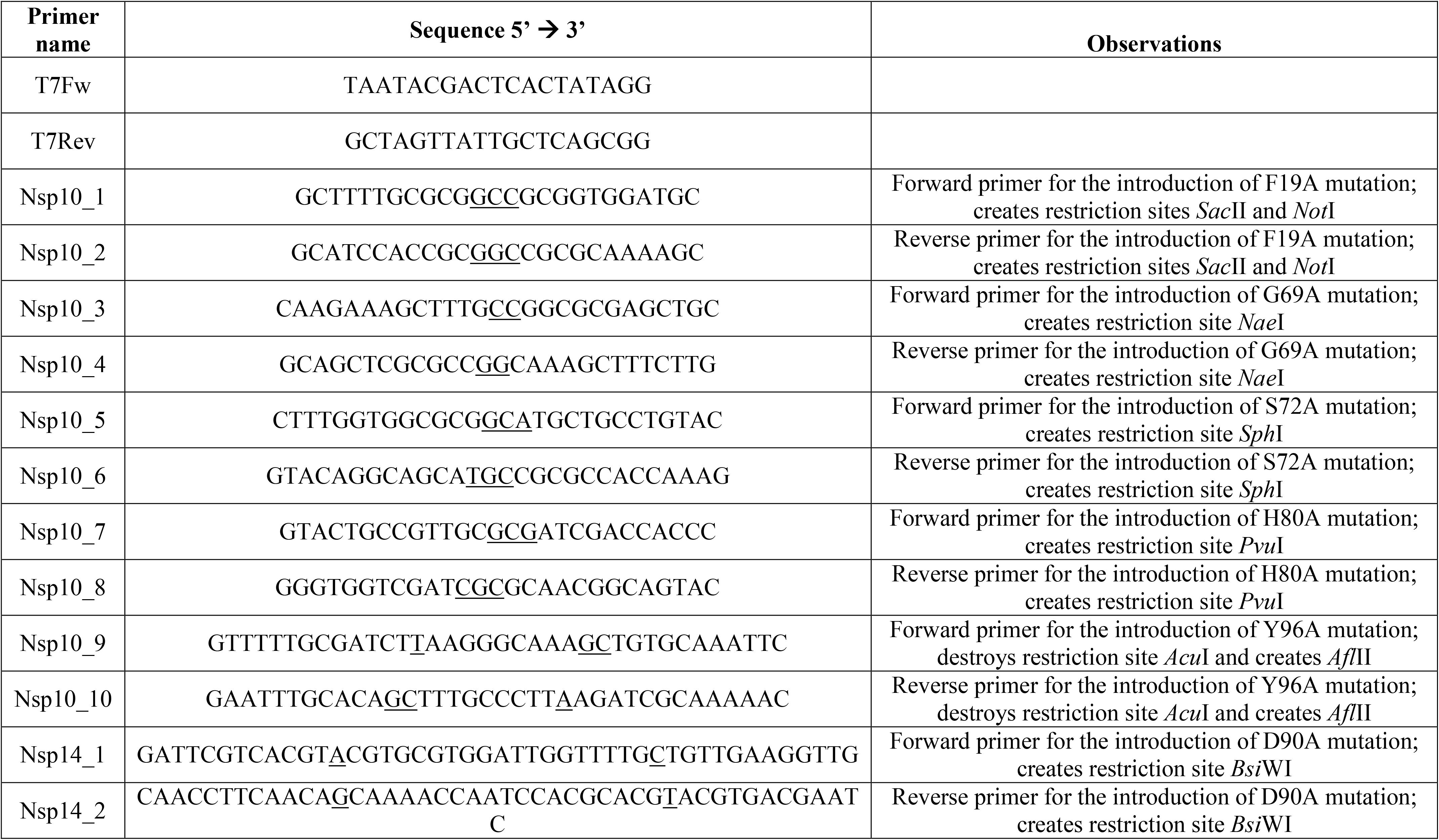
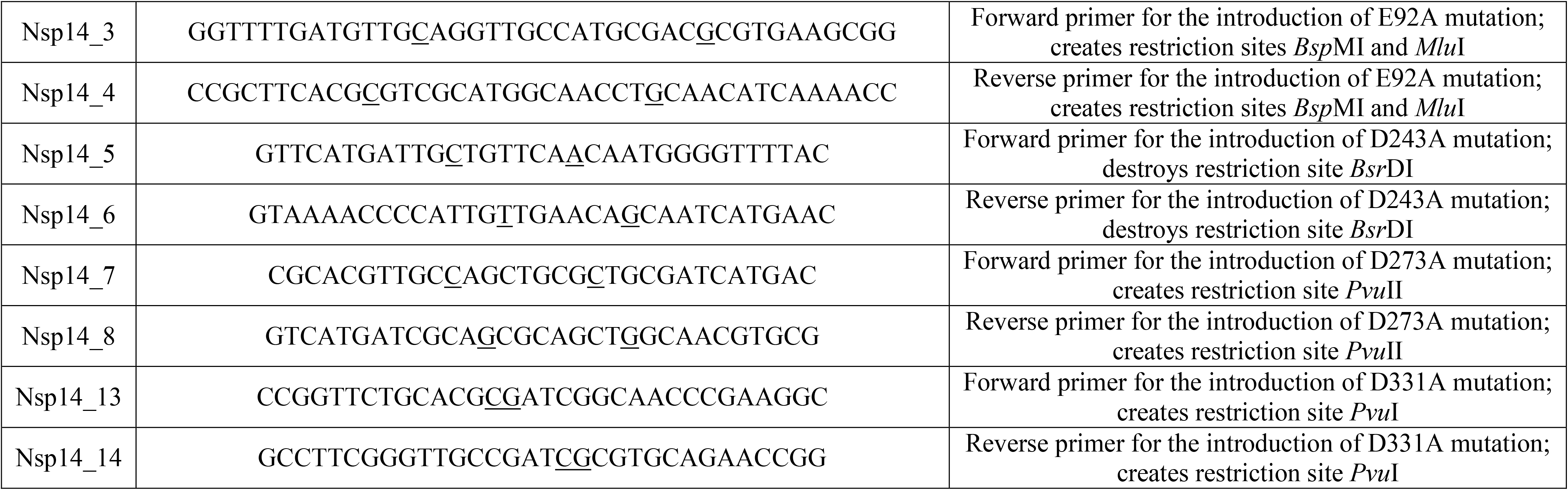
Primers used in this study; bases changes are underlined

**Table S2.**
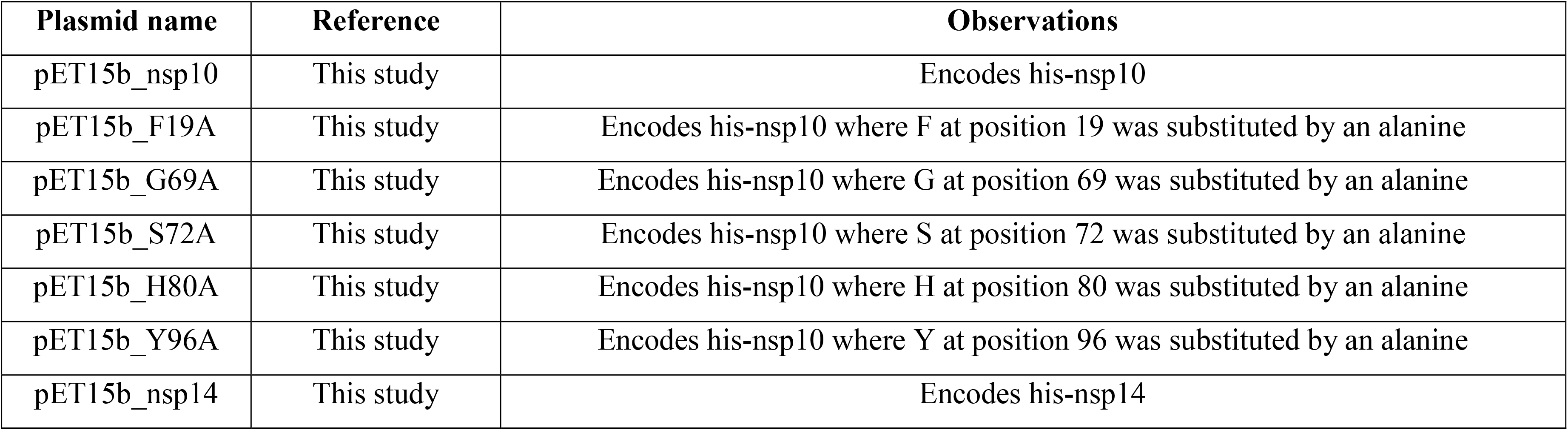
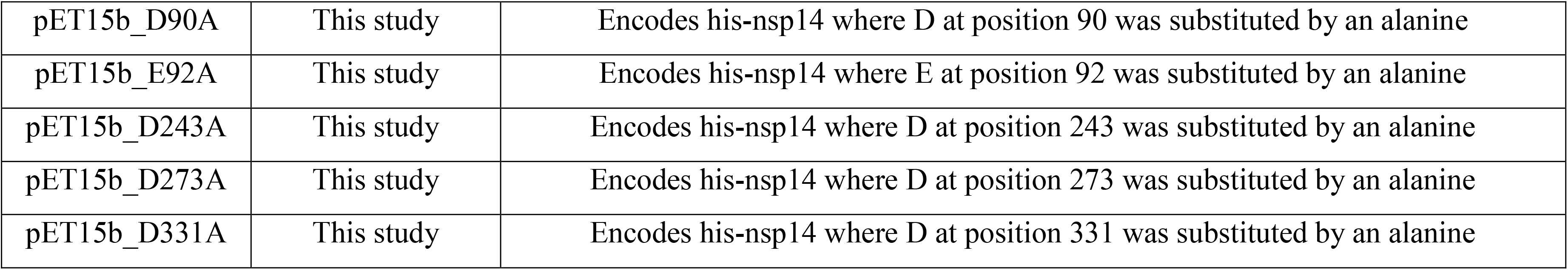
Plasmids used in this study

